# The actin cytoskeleton governs apical mitosis and daughter cell dispersion in intestinal epithelia

**DOI:** 10.1101/183301

**Authors:** Kara L. McKinley, Nico Stuurman, Ophir D. Klein, Ronald D. Vale

## Abstract

Cell proliferation is critical for maintaining the absorptive, protective and regenerative functions of the small intestine throughout adulthood. Interphase nuclei are positioned near the basal surface of the intestinal epithelium, but during mitosis, chromosomes are located apically. The molecular basis for apical-basal DNA positioning and its consequences for tissue homeostasis are poorly understood. Here, we image and pharmacologically perturb these behaviors in live murine intestinal organoids. We find that apical and basal DNA movements occur as a result of mitosis-coupled actin rearrangements that alter the basolateral shape of dividing cells, while the apical cell surface remains confined by cell-cell contacts that persist throughout mitosis. Strikingly, these polarized shape changes allow neighboring cells to insert between nascent daughters, intermingling cells of different lineages. In summary, polarized rearrangements of the actin cytoskeleton govern the mitotic behavior of intestinal epithelial cells and lead to interspersion of cell lineages.

## Introduction

Epithelia undergo extensive cell division to expand and shape tissues during development and to replace old or damaged material in the adult. Since epithelia provide critical barrier functions required for the compartmentalization of multicellular organisms, cell division must occur in a manner that preserves epithelial integrity. Dividing cells must orient the division plane along the appropriate axis of the tissue and maintain cell-cell junctions with their neighbors. In addition, the functions of diverse epithelia rely on the concerted action of multiple cell types, and therefore divisions must generate daughter cells of specific cell fates that are appropriately positioned within the tissue. Thus, epithelial cell division processes must maintain the pattern, shape, and essential functions of the tissue.

A conspicuous behavior during cell proliferation in many epithelia is the occurrence of mitosis on the apical surface of the tissue. During development, apical mitosis occurs in pseudostratified epithelia, densely packed tissues in which nuclei reside at multiple positions along the apical-basal axis, generating the appearance of layered (stratified) cells (reviewed in Norden, 2017). In these tissues, nuclei move apically during the G2 phase of the cell cycle and basally during G1. These movements of interphase nuclei, which cytologically resemble nuclei during interkinesis of meiosis, were coined interkinetic nuclear migration (Sauer, 1935, 1936). The functional significance of interkinetic nuclear migration for epithelia has remained elusive, although it has been proposed to play a role in cell fate decisions (Del Bene et al., 2008), and defects in this process contribute to aberrant development of neural tissues (Doobin et al., 2016; Strzyz et al., 2015).

Interkinetic nuclear migration is observed across numerous pseudostratified epithelia during development (Meyer et al., 2011), but distinct mechanisms underlie this behavior in diverse organs and organisms. In the rodent neuroepithelium, nuclei are transported along microtubules, which are oriented with their minus ends towards the apical surface. In this tissue, apical movement during G2 is achieved by tethering of the motor protein dynein to nuclear pores (Hu et al., 2013; Tsai et al., 2005; Tsai et al., 2010), and basal movement during G1 occurs through movement of the nucleus by kinesins (Tsai et al., 2010). In contrast, in systems including the zebrafish retina and fly wing disc, apical movement during interkinetic nuclear migration requires actin and myosin (Meyer et al., 2011; Norden et al., 2009). However, it remains unclear how nascent nuclei move basally after division in these tissues.

Apical mitosis has also been observed in the adult small intestine (Carroll, 2017; Fleming et al., 2007; Jinguji and Ishikawa, 1992; Trier, 1963), a columnar epithelium in which interphase nuclei localize at approximately the same position along the apical-basal axis, close to the basal surface. However, the molecular mechanisms that underlie apical mitosis in the context of this tissue architecture have not been defined. The small intestinal epithelium is one of the most proliferative tissues in mammals, with most cells turning over in 3-5 days in both mice and humans (Barker, 2014). Divisions of stem cells residing in the crypts of Lieberkühn replenish the stem cell pool and generate absorptive and secretory progenitor cells, which in turn produce differentiated cells that carry out the absorptive and protective functions of the gut (Gracz and Magness, 2014). Absorptive and secretory cells are interspersed between one another along the crypt and villus length (Yang et al., 2001). In addition, stem cells are interspersed with Paneth cells (derived from secretory progenitors) at the crypt base (Snippert et al., 2010), and this alternating arrangement has been proposed as a key architectural element of the stem cell niche (Sato et al., 2011). The cell behaviors that disperse new cells to maintain this intermixed pattern are not understood.

Here, we use adult murine small intestinal organoids to image epithelial mitotic cell behaviors at high temporal and spatial resolution. We find that persistent junctions with neighboring cells cause the apical cell surface to resist shape changes during cell division. In contrast, the basolateral cell surface undergoes dramatic changes during the cell cycle, including rounding at mitotic onset, non-concentric furrow ingression during cytokinesis, and protrusion after division to re-establish the interphase architecture. These polarized, actin-dependent, mitotic cell shape changes are responsible for the observed apical and basal DNA movements. Importantly, these behaviors allow neighboring cells to insert into the cytokinetic furrow of dividing cells, resulting in interspersion of cells of different lineages. Our results support a model in which mitosis-coupled actin remodeling in cells that remain apically confined by cell-cell junctions results in apical mitosis and interspersing of cell lineages during divisions in the intestinal epithelium.

## Results

### Real-time imaging of intestinal organoids reveals that mitotic cells displace apically during rounding

To understand the molecular basis for apical mitosis in the small intestine, we performed time-lapse imaging of murine small intestinal organoids, which have been extensively described to recapitulate aspects of intestinal homeostasis *ex vivo* (Kretzschmar and Clevers, 2016; Sato et al., 2009) (Fig. 1A). Time-lapse imaging of transgenic organoids expressing fluorescent histone H2B to mark DNA confirmed that chromosome segregation occurred on the apical surface of the epithelium (Fig. 1B, C), as observed in fixed sections from mouse and human small intestinal epithelia (Fleming et al., 2007; Jinguji and Ishikawa, 1992; Trier, 1963). Using organoids in which cell membranes were fluorescently labeled we observed that, during mitosis, the basolateral cell boundary retracted toward the apical surface as columnar cells converted to a rounded geometry (Movie S1). To determine the extent to which mitotic cells retained contact with the basal surface while positioned apically, we sought to label the boundary of a subset of cells within the organoid. To achieve this, we used the Cre reporter allele, *R26^mTmG^*, which converts from red membrane fluorescence to green membrane fluorescence upon recombination. We recombined a subset of cells using low levels of induction of the intestinal epithelial Cre driver *Vil1^CreER^*. We observed that mitotic cells were connected to the basal surface of the epithelium by one or more fine membranous processes (Fig. 1D). Thus, both the chromosomes and the majority of the cytoplasm are positioned apically during mitosis in the intestinal epithelium.

**Figure 1.**
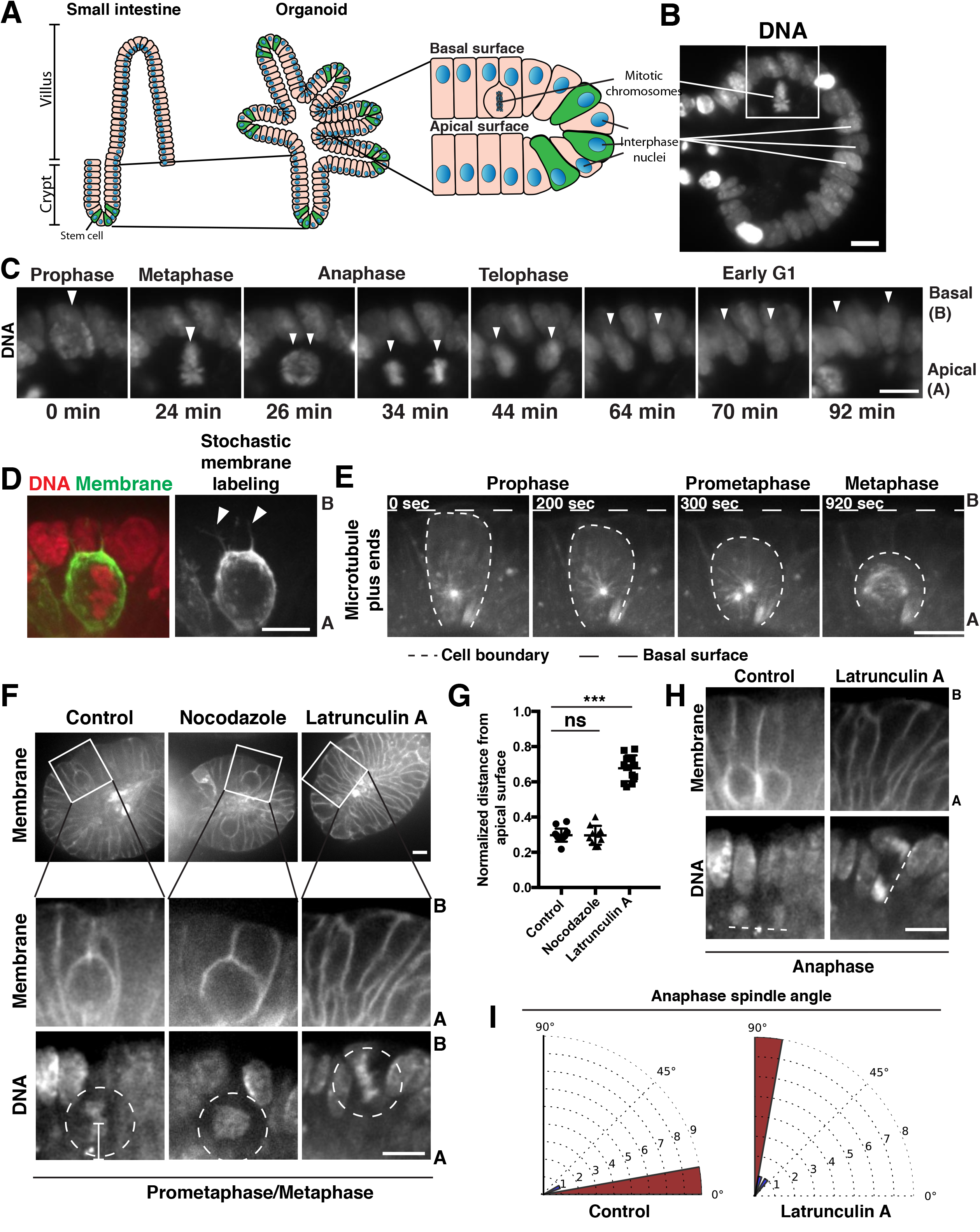
Mitotic rounding underlies apical cell displacement on mitotic entry. A) Cartoon depicting the derivation of organoids from the small intestinal epithelium and the observation of basal positioning of interphase nuclei and apical positioning of mitotic chromosomes, as shown in (B). B) Fluorescent image of DNA (labeled with H2B-mScarlet) in a live organoid. White box indicates mitotic cell highlighted in (C). C) Frames from time-lapse imaging of a mitotic cell (arrowheads) in a live organoid in which DNA is fluorescently labeled with H2B-mScarlet. Images are scaled with ɣ adjustment. Representative of n > 50. D) Fluorescent image of a metaphase cell in a live organoid, in which DNA is fluorescently labeled with H2B-mScarlet. The membranes of a subset of cells within the organoid have been labeled with GFP by inducing low levels of recombination of the *R26^mTmG^* reporter with an inducible, pan-intestinal epithelial Cre *(Vil1^CreER^)*. Arrowheads indicate thin membranous processes that maintain the connection of the mitotic cell to the basal surface. Representative of n > 30, although the number of processes observed per cell is variable. E) Frames from time-lapse imaging of a mitotic cell in a live organoid in which microtubule plus-ends are labeled with EB3-GFP. Representative of n > 20. F) Frames from time-lapse imaging of mitotic cells in live organoids treated with cytoskeletal inhibitors for 30 min before initiation of imaging (nocodazole: 5 µM, latrunculin A: 4 µM). Membranes are labeled with td-Tomato from the *R26^mTmG^* allele in the absence of a Cre driver. DNA is labeled with SiR-DNA dye. Circles indicate mitotic chromosomes; bar indicates measurement of distance from apical surface as quantified in (G). G) Quantification of the distance of mitotic chromosomes from the apical surface of the organoid epithelium following treatment with cytoskeletal inhibitors, normalized to the total apical-basal height of the epithelium, n ≥ 10. ns: not significant; *** p < 0.001, Student's t-test. Cells were treated with inhibitors for 30 min before initiation of imaging, and only cells that entered mitosis during the subsequent 1 h 15 min were quantified. H) Frames from time-lapse imaging of anaphase cells in live organoids treated with cytoskeletal inhibitors as described in (F). Images correspond to later time points of the same cells shown in (F). Dashed lines underline anaphase chromosome masses. I) Quantification of anaphase spindle orientation compared to the plane of the epithelium following vehicle and Latrunculin A treatment, on a 0 - 90° scale, n = 10. Scale bars, 10 µm.

We next analyzed the timing of apical movement relative to mitotic progression. During interkinetic nuclear migration, apical chromosome positioning initiates during G2 phase of the cell cycle (Hu et al., 2013; Leung et al., 2011). In contrast, in the intestinal epithelium, we observed that chromosome movements initiated with chromosome condensation at mitotic entry (Fig. 1C, Movie S2). Imaging of microtubule plus ends using fluorescent End-Binding protein 3 (EB3) revealed that the mitotic spindle assembled concurrently with the rounding and retraction of the dividing cells from the basal surface; ultimately the metaphase spindle assembled in proximity to the apical surface (Fig. 1E, Movie S3). Collectively, these data indicate that apical DNA movements initiate at mitotic entry concurrently with cell rounding to position the chromosomes and mitotic spindle apically.

### Rearrangement of the actin cytoskeleton is required for apical mitotic positioning and planar spindle orientation

As apical chromosome positioning was coincident with rounding of the cell body onto the apical surface, we tested whether mitotic cell rounding was required for apical DNA movements. Dramatic reorganization of the actin cytoskeleton is critical for cells to adopt a spherical geometry during mitosis (reviewed in Lancaster and Baum, 2014; Thery and Bornens, 2008). To test whether mitotic rounding is required for apical DNA displacement, we performed live cell imaging in the presence of the actin depolymerizing small molecule latrunculin A. Latrunculin A (4 µM) abrogated cell rounding, such that, during mitosis, cells remained extended along the full apical-basal axis (Fig. 1F). Strikingly, mitosis occurred on the basal surface of these elongated cells following F-actin depolymerization (Fig. 1F and G, Movie S4). In contrast, cells in which microtubules were depolymerized by nocodazole treatment (5 µM) rounded onto the apical surface during mitosis (Fig. 1F and G, Movie S5) similarly to control cells (Fig. 1F and G, Movie S6). These results indicate that apical displacement in the rodent intestinal epithelium does not require microtubule-motor based transport, as in the rodent neuroepithelium (Hu et al., 2013; Tsai et al., 2005). These data suggest that cell rounding due to remodeling of the actin cytoskeleton is required for apical mitosis in the small intestine.

Mitotic cells in latrunculin-treated organoids also exhibited a severe spindle orientation defect. In contrast to the planar orientation robustly observed during normal divisions in the intestinal epithelium, latrunculin-treated cells underwent orthogonal divisions (Fig. 1H and I, Movie S4, S6). These data indicate that actin-dependent mitotic rounding is required for the planar orientation of the spindle. Collectively, our data suggest that actin-based cell rounding is required for mitotic apical displacement and planar spindle orientation in the intestinal epithelium.

### Mitotic cell shape changes are restricted to the basolateral surface

Although mitotic rounding necessitates a decrease in the height of the cell undergoing mitosis to achieve a spherical geometry, *a priori* such a height decrease could be achieved by retraction of the basal surface, the apical surface, or both surfaces. However, we observed invariant retraction of the basal surface (Fig. 1). Electron microscopy of the intestinal epithelium has revealed that tight junctions are maintained during mitosis in the intestinal epithelium (Jinguji and Ishikawa, 1992). Therefore, an appealing hypothesis is that persistent cell-cell junctions and the resulting apical actomyosin network resist deformation by mitotic rounding forces, thereby requiring that cells round onto the apical surface. Consistent with this, only the basolateral surface of cells adopted a rounded shape during mitosis (Fig. 2A, Fig. S1A). In contrast, the apical surface was not appreciably altered during mitosis. Thus, when viewed from the apical surface, mitotic cells resembled their interphase neighbor cells in terms of the shape and perimeter of the footprint (Fig. 2A and B; Fig S1A), and myosin enrichment (Fig. S1B). These data indicate that the apical surface of the epithelium resists mitotic cell rounding.

**Figure 2.**
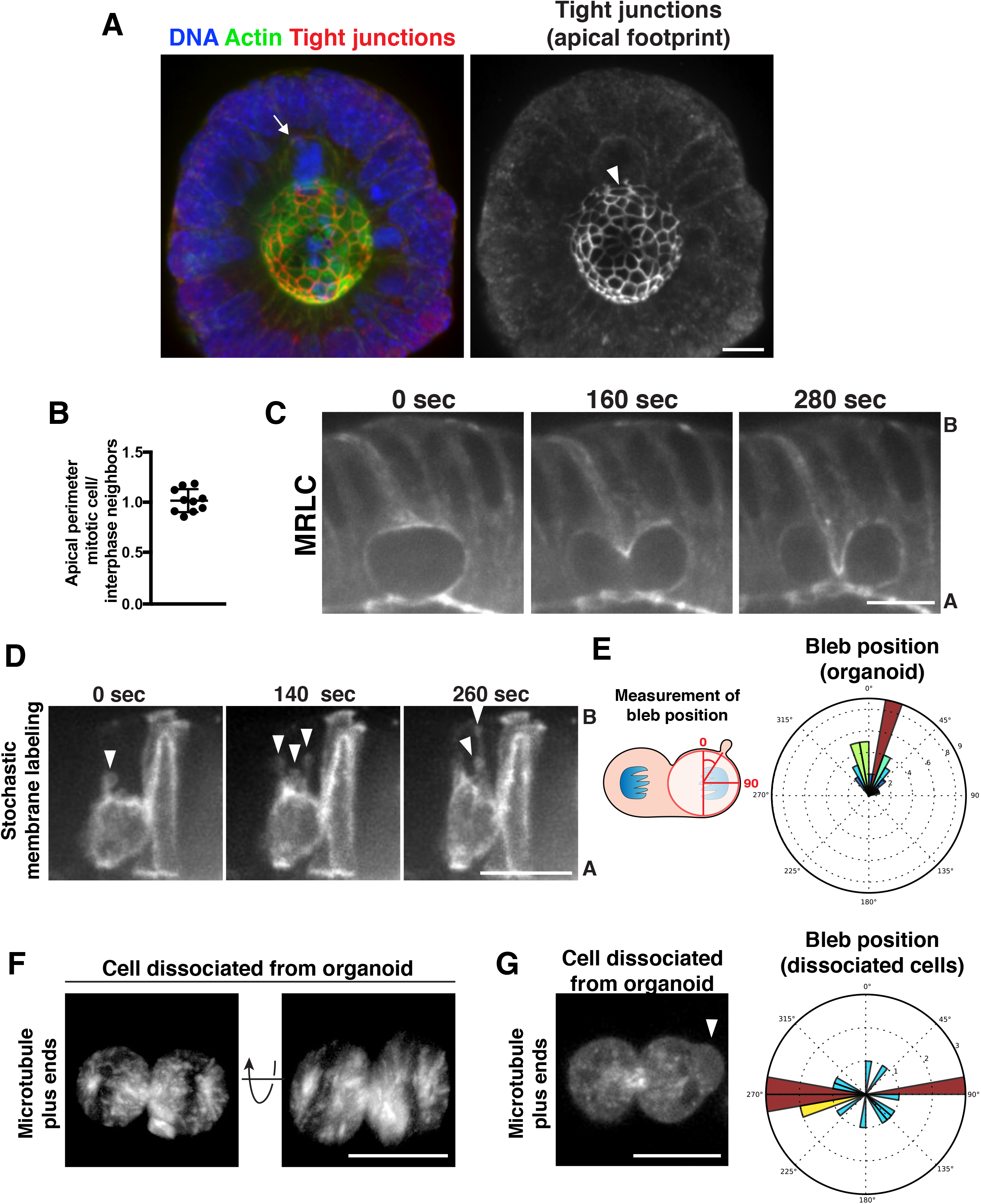
Mitotic cell shape changes are confined to the basolateral surface by contacts with neighboring cells. A) 3 dimensional reconstruction of immunofluorescence images. Arrow indicates mitotic cell. Arrowhead indicates the corresponding apical footprint. DNA was labeled with Hoechst 33342, actin was labeled with Alexa488-phalloidin and tight junctions were labeled with anti-ZO-1. B) Quantification of the perimeter of the apical footprint of mitotic cells compared to interphase cells. The apical footprint was determined from anti-ZO1 immunofluorescence. Each data point represents the ratio between the apical perimeter of a mitotic cell and the average apical perimeter of 4 of its interphase neighbors. n = 10. C) Frames from time-lapse imaging of cytokinesis in a live organoid in which myosin regulatory light chain (MRLC) has been fluorescently labeled with mScarlet. Representative of n > 30. D) Frames from time-lapse imaging of *Vil1^CreER^; R26^mTmG^* organoids induced as in Fig. 1D to stochastically label a subset of cell membranes in the organoid. Arrowheads indicate blebs. Note that the division occurred along the imaging plane, such that the other daughter cell is “behind” the imaged daughter cell. Representative of n > 20. E) Left: Schematic of quantification of bleb position in the organoids. Right: Quantification of bleb position during anaphase and cytokinesis in the organoids. All blebs emerging throughout cytokinesis were quantified across 10 cells. F) 3 dimensional reconstruction of a frame from live imaging of a cell dissociated from EB3-GFP organoids undergoing cytokinesis. A representative cell is rotated towards the viewer. EB3-GFP labeled organoids were used to image furrowing due to poor 3D reconstruction of membrane labels. G) Left: frame from time-lapse imaging of a cell dissociated from EB3-GFP organoids undergoing cytokinesis. Arrowhead points to bleb. Image is scaled with gamma adjustment to visualize cytoplasmic GFP. Right: quantification of bleb position during anaphase and cytokinesis in cells dissociated from organoids, measured as in part (E), n = 8. Scale bars, 10 µm.

The characteristic changes in cell shape that occur upon mitotic exit were also restricted to the basolateral cell surface. Cytokinesis in tissue culture cells involves concentric constriction of a furrow formed around the midzone of anaphase cells. However, time lapse imaging of intestinal organoids labeled with fluorescent myosin regulatory light chain (MRLC) revealed that the cytokinetic ring ingressed from the basolateral surfaces until it reached the apical surface (Fig. 2C, Movie S7, S8; also see Fleming et al., 2007). In contrast, the apical surface did not exhibit any apparent furrowing (Fig. 2C, Movie S7, S8). Additionally, we observed dramatic polarized membrane blebbing accompanying cytokinesis (Fig 2D, E, Movie S9). In tissue culture, blebbing occurs at the poles due to local cortical disruptions downstream of chromatin-derived signals (Kiyomitsu and Cheeseman, 2013; Rodrigues et al., 2015). In contrast, blebbing in the intestinal epithelium occurred only from the basal surface of dividing cells (Fig. 2D, E, Movie S9). Together, these data suggest that changes in cell shape during mitotic progression are confined to the basolateral surface, while the apical surface maintains a fixed architecture.

As mechanical coupling between cells in the epithelium is achieved through apical cell junctions and the resulting contractile actomyosin network, we sought to test the role of cell junctions in rendering the apical surface refractory to cell shape changes. To assess this, we dissociated organoids into single cells or pairs of cells and performed time-lapse imaging of mitotic exit. In contrast to the non-concentric cytokinesis and basal blebbing observed in the intact tissue (Fig. 2C, D, E), in dissociated cells we observed concentric furrows (12/15 cells, Fig. 2F, Movie S10) and polar blebbing (Fig. 2G), as observed for tissue culture cells. Taken together, these data indicate that cell-cell junctions maintain a persistent apical architecture of the intestinal epithelium, thereby polarizing mitotic cell shape changes.

### Actin-based cell elongation drives reinsertion of nascent nuclei onto the basal surface after mitotic exit

Following cytokinesis, we observed that the basal edge of nascent daughters formed a protrusive front that extended basally to re-establish contact with the basal surface (Fig. 3A, Movie S11). The protrusion of this front resembled protrusion of the leading edge of migrating cells, a process in which the actin cytoskeleton plays a central role (Ridley, 2011). Therefore, we next tested the requirement for actin in basal reinsertion. As actin disruption blocks the initial displacement of mitotic cells to the apical surface (Fig. 1F, G), determining the requirements for actin in basal reinsertion required that mitotic cells be positioned on the apical surface before disrupting actin. To achieve this, we first blocked cells on the apical surface by arresting them in mitosis with the mitotic kinesin (Eg5) inhibitor S-trityl-L-cysteine (STLC, 10 µM). Mitotically arrested cells did not re-insert onto the basal surface over at least three hours (Fig. S2A, B, C), indicating that mitotic exit is required for basal reinsertion. To induce exit from mitosis in the presence of STLC, we added compounds that inhibited either the spindle assembly checkpoint (SAC; Mps1 inhibitor, 2 µM) or cyclin-dependent kinase (CDK, RO-3306, 10 µM). In both cases, we found that these 4N cells reinserted onto the basal surface in a manner similar to untreated cells (Fig. S2A, B, C, Movie S12). Thus, mitotic exit and reversal of CDK phosphorylation are sufficient for basal reinsertion, even in the absence of chromosome segregation.

**Figure 3.**
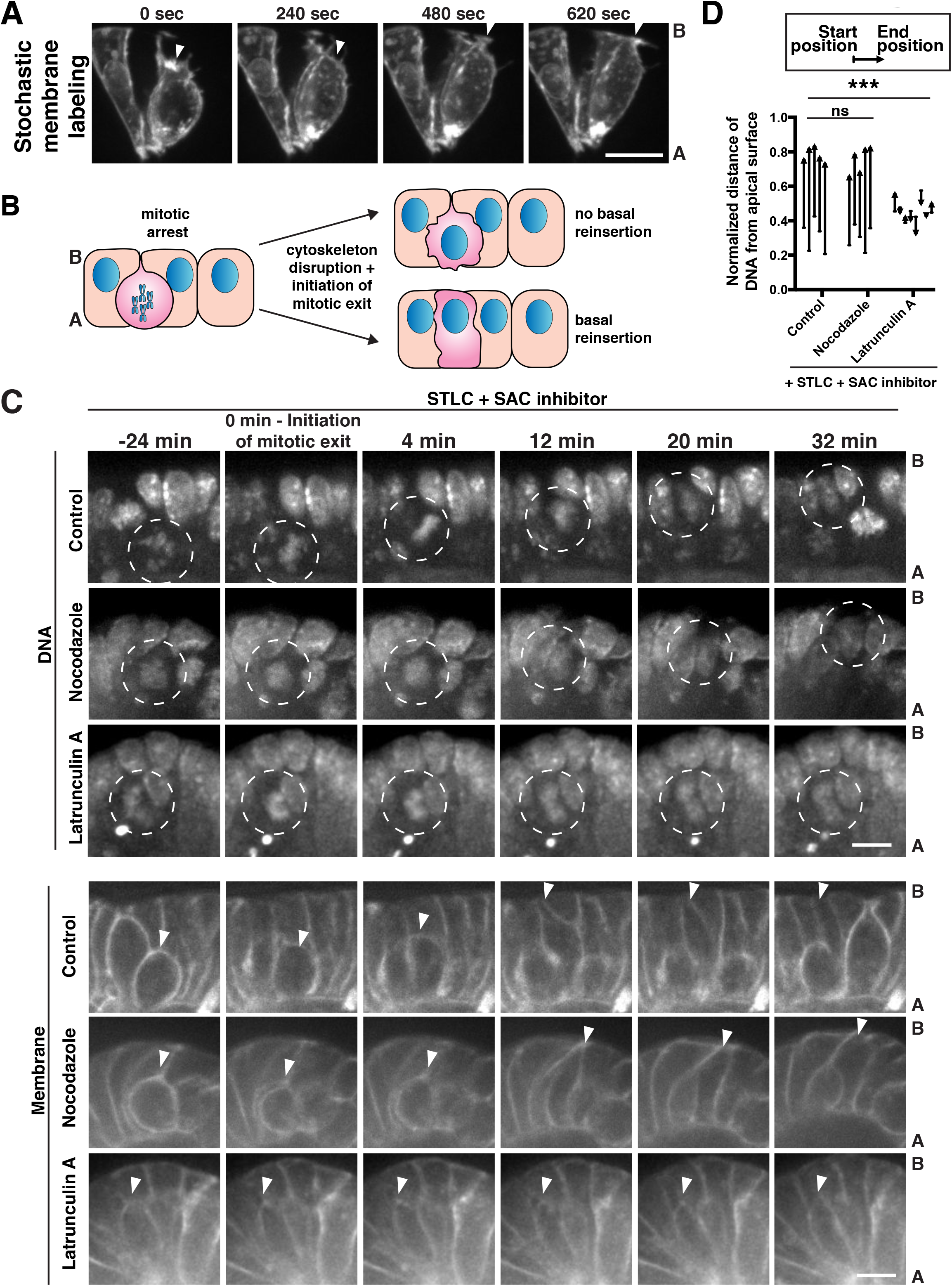
Actin-based cell elongation drives reinsertion of nuclei onto the basal surface following mitotic exit. A) Frames from time-lapse imaging of *Vil1^CreER^; R26^mTmG^* organoids induced as in Fig. 1D to stochastically label a subset of cell membranes in the organoid. Arrowheads indicate protrusive cell front. B) Cartoon illustrating the strategy to analyze chromosome movements in organoids following mitotic arrest with S-trityl-L-cysteine (STLC, 10 µM), pharmacological disruption of the cytoskeleton, and induction of mitotic exit with the SAC inhibitor AZ3146 (2 µM). C) Frames from time-lapse imaging of live organoids testing the cytoskeletal requirements for basal reinsertion as described in (B) and Fig. S2D. DNA was labeled with SiR-DNA dye (1 µM), membrane was labeled with the *R26^mTmG^* allele in the absence of recombination. Circles indicate chromosomes; arrowheads indicate the basal edge of the corresponding cell membrane. D) Quantification of DNA position before the addition of SAC inhibitor AZ3146 (starting position), and at chromosome decondensation (end position) as depicted in (C). Each arrow corresponds to the position of the DNA from one cell, before and after the treatment, normalized to the total apical-basal distance of the epithelium. Arrowheads point towards the end position after mitotic exit. n ≥ 5, ns: not significant, ***: p < 0.001, Student's t-test of distances moved (end position - start position). Scale bars, 10 µm.

Using this mitotic arrest and exit protocol, we were able to test the requirements for the actin and microtubule cytoskeleton for basal reinsertion. Following mitotic arrest with STLC, cells were treated with cytoskeletal inhibitors and mitotic exit was induced using SAC inhibition (Fig. 3B, S2D). Cells were then analyzed by time-lapse microscopy to determine whether they could undergo basal reinsertion. Depolymerization of microtubules with nocodazole before and after mitotic exit did not interfere with the ability of nuclei or the cell boundary to reach the basal surface (Fig. 3C, D, Movie S13). In contrast, disruption of F-actin polymerization with latrunculin blocked basal reinsertion. Instead, the nucleus reformed its interphase morphology on the apical surface and the cell boundary did not protrude toward the basal surface (Fig. 3C, D, Movie S14). Collectively, these data indicate that the actin cytoskeleton drives basal reinsertion of daughter cells after mitotic exit.

Since actin also plays a critical role in cytokinesis, we sought to test whether basal reinsertion depends upon successful completion of cytokinesis. To inhibit cytokinesis without disrupting actin, we utilized the Polo-like kinase 1 (Plk1) inhibitor BI2536 (Lenart et al., 2007; Steegmaier et al., 2007) (10 µM), since blebbistatin, a myosin II inhibitor, did not disrupt cytokinesis at the limits of its solubility in this system (data not shown). Although Plk1 is required for proper spindle assembly during mitosis, inhibition of Plk1 after metaphase allows cells to undergo chromosome segregation without cytokinesis (Burkard et al., 2007; Petronczki et al., 2007). We found that in the absence of cytokinesis, nuclei reinserted normally onto the basal surface (Fig. S2E, F), indicating that cytokinesis is dispensable for basal movement.

### Apical mitosis and non-concentric cytokinesis permit insertion of neighbors between daughter cells

Finally, we investigated the consequences of the observed mitotic cell shape changes for the architecture of the epithelium. For these experiments, we utilized organoids in which the cytoplasm of cells of the secretory lineage was labeled with RFP (*Atoh1^CreER^; R26^Rfp^)*. Cells of the secretory lineage are frequently interspersed with small patches of non-secretory cells, both in organoids and *in vivo* (Fig. 4A) (Yang et al., 2001). Therefore, we could directly visualize the position of secretory daughters relative to non-secretory neighboring cells. Strikingly, we observed that the two secretory daughter cells frequently became separated from one another by non-secretory cells as they reinserted toward the basal surface (31/50 divisions, Fig. 4A, Movie S15). To determine the extent of daughter cell contact in all dimensions, we analyzed secretory lineage-labeled organoids in three dimensions over time using light sheet microscopy (single plane illumination microscopy - SPIM) (Wu et al., 2013). These analyses revealed two categories of daughter cell geometry after basal reinsertion: either the daughters maintained contact along the full apical-basal cell length (5/10 daughter pairs), or the daughters adopted a V-shaped geometry, in which they remained closely apposed on the apical surface, but had no apparent contact on the basal surface (Fig. 4B, Movie S16, 5/10 daughter pairs). Thus, a subset of daughter cells separated from one another as they reinserted toward the basal surface after apical mitosis, resulting in an interspersing of cell lineages.

**Figure 4.**
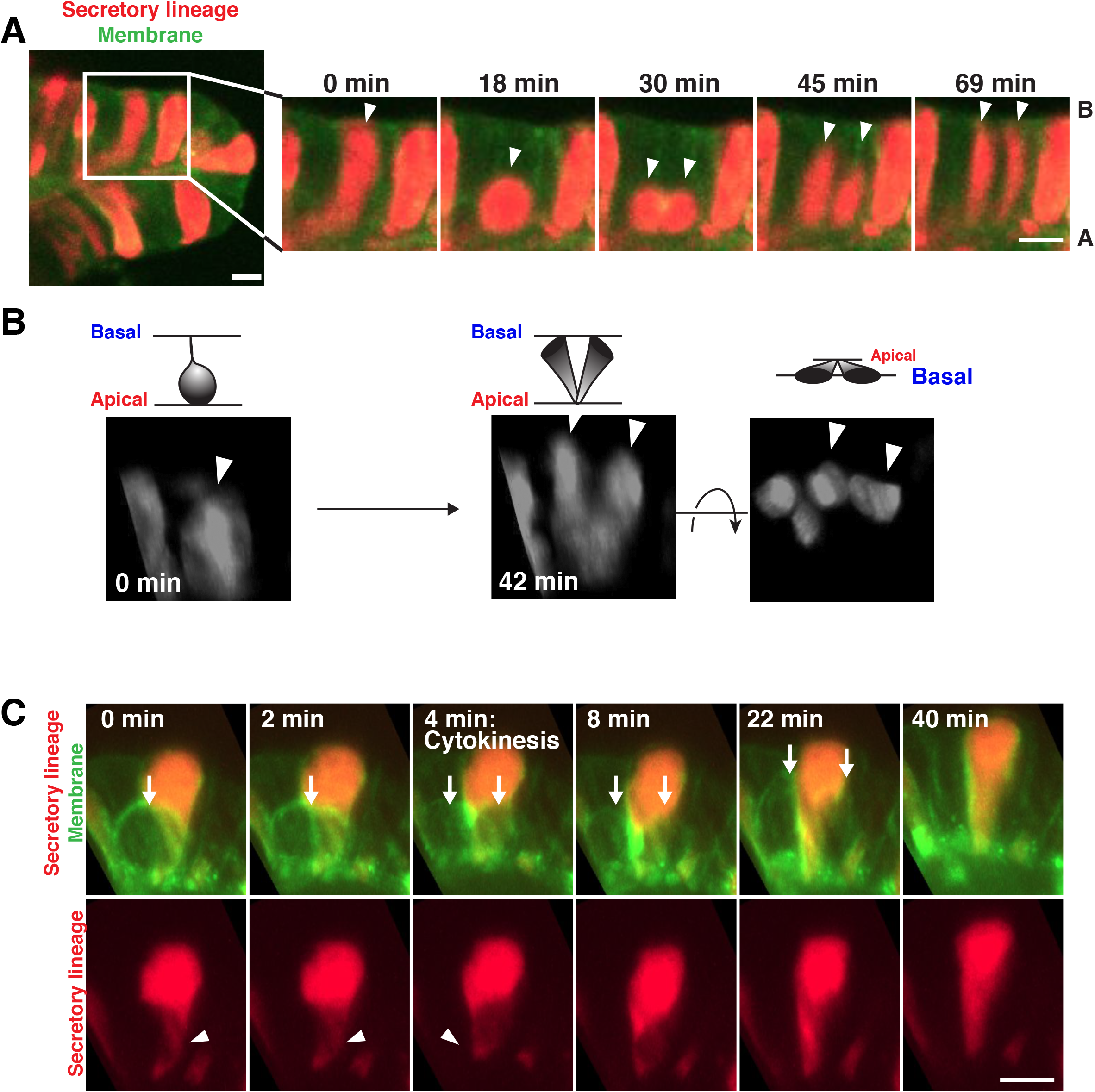
Neighboring cells insert into the cytokinetic furrow to displace reinserting daughter cells. A) Frames from time-lapse imaging of a dividing cell of the secretory lineage (red, *Atoh1^CreER^; R26^rfp^*). Arrowheads indicate dividing cell and nascent daughters. B) Frames from 3D reconstructed time-lapse SPIM imaging of a dividing cell of the secretory lineage. Left panel shows mitotic cell before division, right panels show two views of the resulting daughter cells after basal reinsertion. Arrowheads indicate dividing cell and nascent daughters. C) Frames from 3D reconstructed time-lapse SPIM imaging of a cell of the secretory lineage (red) inserting in the cytokinetic furrow of a dividing cell (green). Arrows indicate the dividing cell. Arrowheads indicate the cytoplasmic process of the inserting neighboring cell. Top panels show both the dividing and neighboring cell; bottom panels isolate the neighboring cell. Scale bars, 10 µm.

We next sought to understand the origins of the V-shaped daughter cell geometry. As described above, we observed two behaviors that could potentially contribute to daughter cell separation following division. First, daughter cells underwent non-concentric cytokinesis (Fig. 2C). Second, newly divided daughters extended a protrusive front to re-establish contact with the basal surface (Fig. 3A). Therefore, neighboring cells could intercalate between the two daughters by inserting into the position generated by the ingressing cytokinetic furrow. Alternatively, daughter cells could separate after cytokinesis, as they extend their protrusive fronts towards the basal surface. Time-lapse light sheet microscopy and 3-dimensional reconstruction revealed that neighboring cell cytoplasm inserted into the furrow during cytokinesis, with daughter cells subsequently elongating around this insertion (Fig. 4C, Movie S17). Thus, apical mitosis and nonconcentric cytokinesis permit insertion of neighboring cells into the cytokinetic furrow, and the inserted neighbor displaces the daughter cells from one another as they re-insert on the basal surface.

## Discussion

A central question in developmental biology is how cell rearrangements within epithelia give rise to complex structures. This challenge continues after embryonic development into adulthood, as new cells must be produced and distributed to replenish old or damaged material. During homeostasis of the adult small intestine, interphase nuclei are positioned basally, but mitotic chromosomes are observed on the apical surface of the epithelium (Jinguji and Ishikawa, 1992; Trier, 1963). We found that apical and basal DNA movements were driven by cell cycle-dependent rearrangements of the actin cytoskeleton. These rearrangements altered basolateral cell shape, while the apical surface of mitotic cells resisted shape changes due to cell junctions. These polarized changes in shape allowed neighboring cells to insert between dividing cells, resulting in interspersing of cells of different lineages (Fig. 5).

**Figure 5.**
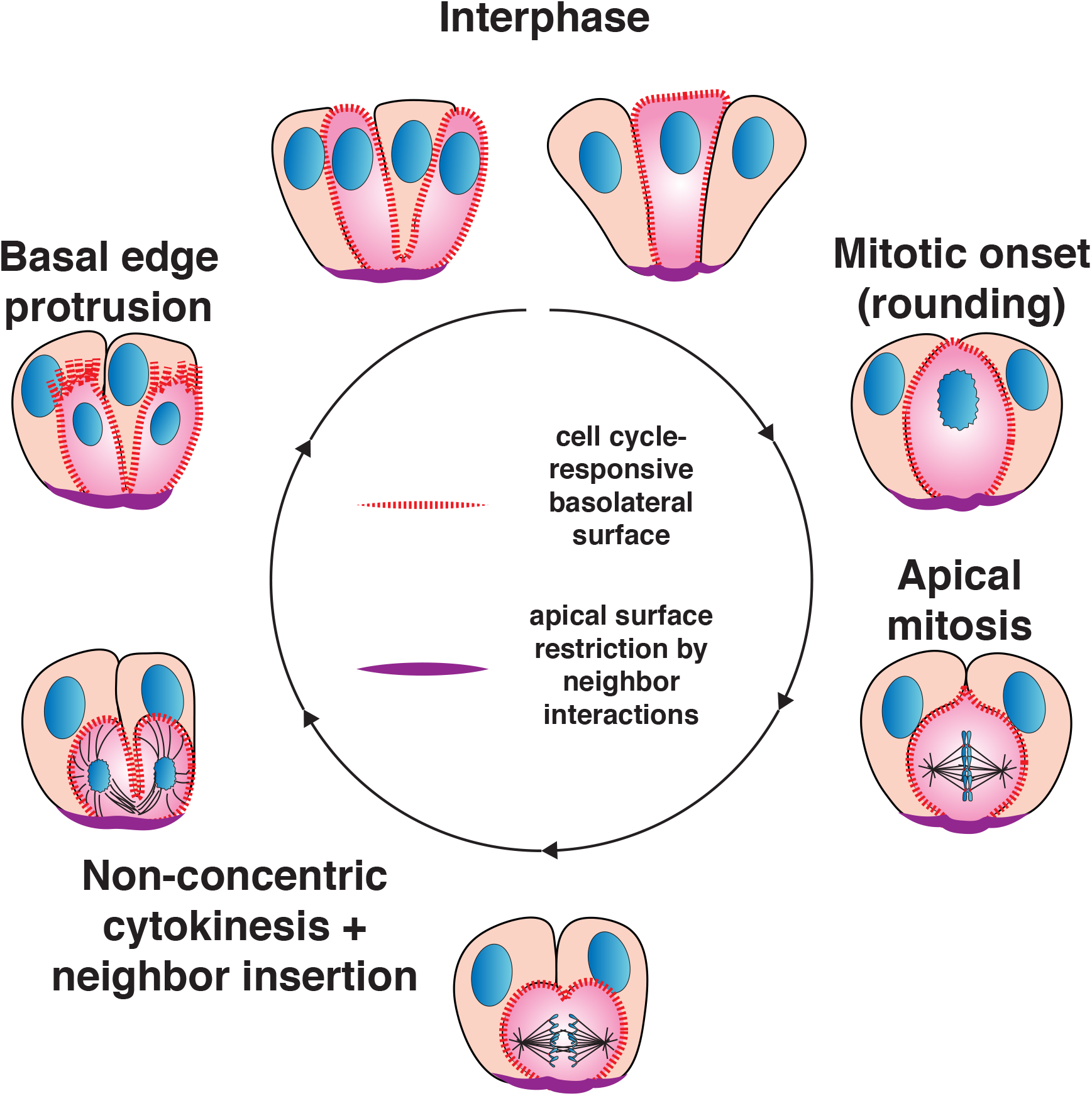
Model for apical and basal mitotic movements and daughter cell separation. The basolateral surface of a cell undergoing mitosis (pink) undergoes dramatic actin-dependent changes in shape (dashed red lines). The apical surface resists mitotic cell shape changes due to contacts with neighboring cells and the persistent actomyosin network (purple lines). Polarization of cell shape changes results in apical and basal movements of the chromosomes and cell body. Neighboring cells can position within the ingressing cytokinetic furrow, displacing daughter cells from one another as they reinsert onto the basal surface.

### Polarized actin rearrangements drive apical and basal positioning of the DNA

Apical mitosis has been best studied in the context of interkinetic nuclear migration in pseudostratified epithelia during development, which involves movement of nuclei along the cell length during interphase. In contrast, our data indicate that apical and basal DNA movements that occur during intestinal homeostasis are a consequence of combining the changes in cell shape required to generate the optimal geometry for mitosis with the fixed apical footprint that maintains the function of the intestinal epithelium as a chemical and mechanical barrier.

Most cells in tissues and in culture have been observed to adopt a spherical geometry that requires dramatic reorganization of the actin cytoskeleton (reviewed in Thery and Bornens, 2008). Conversion of the elongated interphase architecture of intestinal epithelial cells into a sphere requires that cells decrease in height along their apical-basal axis. However, mitotic cells must also maintain their apical tight junctions to ensure ongoing barrier function (Jinguji and Ishikawa, 1992), so the decrease in cell height cannot be achieved by detachment from the apical surface. Rather, mitotic rounding causes cells to almost fully detach from the basal surface and adopt a rounded shape on the apical surface. As a result, the cell body and with it, the assembling mitotic spindle and chromosomes, displace to the apical surface. Mitotic rounding also contributes to apical movements associated with interkinetic nuclear migration in some tissues (Meyer et al., 2011; Spear and Erickson, 2012).

In the intestinal epithelium, we find that failure to achieve this rounded geometry results in orthogonal rather than planar orientation of the mitotic spindle. Recent work has similarly demonstrated that disruption of mitotic rounding impairs both the orthogonal divisions that stratify the murine epidermis and planar divisions in the Drosophila wing disc (Luxenburg et al., 2011; Nakajima et al., 2013). In the murine epidermis, mitotic rounding is required for the correct localization of the protein LGN, which plays a critical role in connecting astral microtubules to the cortex to orient and position the spindle (Luxenburg et al., 2011). In the intestine, a similar interplay between the intrinsic spindle orientation mechanisms and the mitotic rounding apparatus may be at play. Additionally, an appealing model is that the cellular geometry of the intestine may physically impede planar spindle orientation in the absence of mitotic rounding. In the intestinal epithelium, cells are tightly packed together, such that the interphase cell width is smaller than the mitotic spindle length. Thus, achieving planar spindle orientation requires an increase in cell width. Mitotic rounding is driven by intracellular pressure that exerts force (Stewart et al., 2011), which is sufficient to deform neighboring cells (Fig. 4C), allowing the mitotic cell to expand its planar axis to accommodate a planar spindle.

Following apical mitosis, nuclei move basally in both pseudostratified epithelia and the intestinal epithelium. However, the mechanisms that drive these basal movements remain poorly understood outside of the microtubule-based mechanisms in the rodent neuroepithelium (Tsai et al., 2010). In the intestinal epithelium, basal rearrangements after mitosis are particularly critical because apical rounding severely reduces the basal footprint of the cell. We found that actin-dependent extension of the basal edge of the daughter cells reestablishes the interphase basal footprint and basal nuclear positioning following mitosis. We observed that the basal surface of the cell extends a protrusive “front”, resembling a lamellipodium found at the leading edge of many migrating cells (Svitkina, 2013). In contrast, the apical surface maintains its contractile actomyosin network, resembling the retracting rear of migrating cells (Cramer, 2013). Thus, the polarized cytoskeletal architecture that establishes the interphase organization of the intestinal epithelial cell resembles the polarized architecture found in many migratory cells.

### Mitotic cell shape changes are broadly polarized by apical cell-cell junctions

Our data place the cell shape changes that drive apical and basal DNA movements among a suite of polarized cell division behaviors in the intestinal epithelium. Several lines of evidence indicate that these polarized behaviors collectively arise because cell-cell junctions and the ensuing actomyosin network on the apical surface of the epithelium oppose the forces that underlie mitotic shape changes. We observed that the apical footprint of mitotic cells did not differ discernibly from that of interphase cells, indicating that the balance of forces between a cell and its neighbors on the apical surface is maintained when the cell enters mitosis. Following mitosis, we observed non-concentric cytokinesis, in which the furrow ingresses from the basolateral surfaces but without significant furrowing of the apical surface. Non-concentric cytokinesis is extensively observed in metazoan epithelia (Fleming et al., 2007; Founounou et al., 2013; Guillot and Lecuit, 2013; Herszterg et al., 2013; Morais-de-Sa and Sunkel, 2013; Reinsch and Karsenti, 1994), in contrast to concentric cytokinesis observed for cells in culture, unicellular organisms, and during early embryonic development in some organisms. In Drosophila, apically localized adherens junctions are required for non-concentric furrowing (Guillot and Lecuit, 2013; Morais-de-Sa and Sunkel, 2013). Although vertebrates differ significantly from Drosophila in the architecture and composition of their cell-cell junctions, our data suggest that maintenance of cell junctions also causes cytokinesis to proceed non-concentrically in the intestinal epithelium. Similarly, we find that cell junctions polarize blebbing behavior, which occurred only from the basal surface of dividing cells in the intestinal epithelium. Since blebbing is associated with cortical softening (Sedzinski et al., 2011), the polarized blebbing behavior that we observe likely reflects a dramatic mechanical asymmetry in the cortex, with the basolateral surface being mechanically weaker than the apical surface. Together, our data support a model in which persistent apical cell junctions and the resulting contractile actomyosin network broadly oppose cell shape changes during cell division to result in polarized behaviors.

### Consequences of apical mitosis during intestinal homeostasis

An outstanding question for studies of apical mitosis is how this behavior contributes to tissue architecture. Our data reveal two functional consequences of this behavior. First, we find that apical mitosis driven by mitotic rounding is required for planar spindle orientation. This planar spindle orientation is critical to maintain the architecture of the small intestinal epithelium, as orthogonal divisions would drive stratification (Lechler and Fuchs, 2005). Indeed, orthogonal spindle angles are observed in APC mutant intestines and have been proposed to contribute to tumorigenesis (Caldwell et al., 2007; Fleming et al., 2009).

Second, following mitosis, we observed that daughter cells separated from one another as they elongated to the basal surface. Daughter separation after mitosis in the intestinal epithelium was also recently reported by Carroll et al. (Carroll, 2017). Lineage tracing in fixed tissues has long established that cell lineages are interspersed in the intestinal epithelium, with stem cells and Paneth cells adopting a checkerboard pattern at the crypt base (Snippert et al., 2010), and secretory cells interspersed with absorptive cells throughout the epithelium (Yang et al., 2001). Indeed, proximity between Paneth cells and stem cells may play a role in stem cell renewal due to niche-like functions provided by the Paneth cells (Sato et al., 2011). However, it was unclear how this intermingling is achieved. In some systems, such as primordial Drosophila appendages (Gibson et al., 2006), the vast majority of divisions result in daughter cells that share a common interface, as underlined by the employment of contiguous twin spots in Drosophila lineage analyses (Bryant, 1970; Bryant and Schneiderman, 1969). In contrast, evidence has recently emerged for cell division-coupled rearrangements during vertebrate development (Firmino et al., 2016; Higashi et al., 2016; Lau et al., 2015; Packard et al., 2013). Our work extends the paradigm of cell division-coupled rearrangements to the maintenance of cell type patterning during homeostasis of the adult intestinal epithelium.

We propose that the interspersion of cell lineages that we observe accompanying the division can be explained by the polarized cortical responses described above (Fig. 5). In the columnar epithelium of the small intestine, the rounding-based apical displacement dramatically reduces the basal footprint of the cell, leaving behind only thin membranous processes. The cells then undergo non-concentric cytokinesis, and following the division elongate toward the basal surface to re-establish the footprint. We find that the initiating event of daughter separation is the insertion of a neighboring cell between the nascent daughter cells during cytokinesis, thereby displacing the daughter cells from one another when they elongate towards the basal surface. Based on our observation that basal daughter separation is a frequent but not universal event, we speculate that occupation of the cytokinetic furrow by a neighboring cell is opportunistic, arising from relative positioning of neighbor and daughter cells.

### A model for polarized cell cycle behaviors in the intestinal epithelium

Taken together, our work presents a model for cell behaviors and rearrangements arising from polarization of cell shape changes in the intestinal epithelium (Fig. 5). While the apical surface maintains its mechanical properties through cell division due to the persistence of cell-cell junctions, the basal surface undergoes a cycle of cytoskeletal changes that are characteristic of many mitotic cells – rounding at mitotic onset, furrow ingression at cytokinesis and elongation at mitotic exit. As a result, these cytoskeletal changes, which occur symmetrically in isolated cells, become polarized. Thus, we propose that the observed mitotic behaviors in the intestinal epithelium, such as apical mitosis and the interspersion of cell lineages, arise from differences in the mechanical properties of the apical and basal surfaces during the cell cycle. In the future, it will be informative to examine the extent to which these mitotic outcomes of apical confinement, including the insertion of a neighboring cell between daughters, are observed across mammalian epithelia, or are specific to the tight cell packing and high degree of curvature that achieves the cup-shaped geometry of the small intestinal crypt.

## Acknowledgments

We thank members of the Vale and Klein laboratories for technical support, reagents, and critical consideration of the manuscript. We thank Frederic de Sauvage (Genentech) for the Lgr5^DTR-GFP^ allele. We thank Ilia Koev (Biogene) for his gift of a piece of Teflon AF 2400. RDV and NS were supported by the Howard Hughes Medical Institute. This research was funded in part by the Intestinal Stem Cell Consortium – a collaborative research project funded by the National Institute of Diabetes and Digestive and Kidney Diseases (NIDDK) and the National Institute of Allergy and Infectious Diseases (NIAID) (U01DK103147 to ODK). KLM is a Damon Runyon Fellow supported by the Damon Runyon Cancer Research Foundation (DRG-2282-17).

## Materials and Methods

### Mouse lines

Adult mice of the following lines were used to generate organoids.

*R26^mTmG/mTmG^* (Muzumdar et al., 2007) (female)

*Villin^Cre-ERT2/+^* (el Marjou et al., 2004); *R26^mTmG/+^* (male)

*Atoh1^CreERT/+^* (Chow et al., 2006); *R26^rfp/+^* (Madisen et al., 2010); *Lgr5^GFP-IRES-DTR/+^* (Tian et al., 2011) (female)

The strains of these mice were the same as previously described in their references at the time of acquisition but were maintained on mixed backgrounds after breeding between different lines. All experiments involving mice were approved by the Institutional Animal Care and Use Committee of the University of California, San Francisco (protocol #AN151723).

### Organoid preparation, dissociation and immunofluorescence

Small intestinal crypts were isolated from mice and cultured in medium supplemented with human recombinant EGF, human recombinant Noggin and R-Spondin conditioned medium (ENR medium) as described (Sato et al., 2009). Catalog numbers for culture medium components are described in (Mahe et al., 2013). Where indicated, organoids expressing fluorescent proteins were generated by lentiviral transduction as described (Koo et al., 2011). For propagation, organoids were grown in 24-well plastic plates. For spinning disc imaging and immunofluorescence, organoids were grown in 96-well glass bottom dishes (Matriplate, Brooks). For diSPIM imaging, organoids were grown on glass coverslips which were then transferred to the diSPIM imaging chamber. For immunofluorescence, organoids were fixed in 4% PFA in PBS for 30 min before blocking in 3% BSA, TBS, 0.1% Triton X-100. Primary antibody was incubated overnight at 4 degrees and secondary antibody was incubated for > 2 hours at RT. ZO-1 was stained with rabbit anti-ZO-1 (Thermo Fisher RRID:AB_2533456), actin was stained with Alexa 488-Phalloidin (Thermo Fisher Cat # A12379) and DNA was stained with Hoechst 33342 (Molecular Probes H3570).

For organoid dissociation, organoids in one well of a 24 well plate were washed once in PBS before Matrigel was manually disrupted by pipetting in TrypLE Select (Life Technologies) in the well. The plate was then incubated at 37 °C for 7-8 minutes before additional disruption with a P200 pipette. The cell suspension was centrifuged in medium + 5% fetal bovine serum at 1000 x g for 5 min. The pellet was resuspended in Matrigel, allowed to polymerize for 10 min and covered with ENR medium and immediately transferred to the microscope for imaging for 45 min - 1 h.

### Microscopy

Unless otherwise stated, images were acquired on a Yokogawa CSU-X1 spinning disk confocal attached to an inverted Nikon TI microscope, an Andor iXon Ultra 897 EM-CCD camera, using Micro-Manager software (Edelstein et al., 2010). Imaging of 12 x 1 µm z-stacks was performed either at 4-min time intervals with a 40X 1.30 NA Plan Fluor oil objective or a 20X 0.75 NA objective, or at 20 sec time intervals with a 60XA 1.20 NA Plan Apo water immersion objective. Maximum intensity projections of 1-5 Z-stacks are shown unless otherwise noted. 4-dimensional imaging was performed on an ASI diSPIM microscope equipped with 40X 0.80W NA NIR-Apo water dipping objectives, Hamamatsu Flash 4.0 cameras, and 488nm and 561nm solid state lasers from Vortran, using a nightly build of the Micro-Manager software. Temperature was maintained using 3 x 50 ohm resistors attached to the stainless steel incubation chamber holding the coverslip and medium, a 10 kOhm thermistor inserted in the medium and a temperature controller (TE Technology, Inc. TC-48-20). O_2_ and CO_2_ tensions in the medium were kept constant by flowing humidified gas underneath the sample chamber. To allow gas exchange, the sample was placed on a sandwich of 2 x 24x50mm coverslip glasses in which 2 ~12x12 mm windows had been laser-cut and between which a piece of ~37.5 µm thick Teflon AF-2400 (a gift from BioGeneral, Inc.) was placed. Evaporation was minimized by layering mineral oil (Howard) over the sample. 3D reconstructions were generated using a Micro-Manager plugin (https://github.com/nicost/MMClearVolumePlugin) that uses the ClearVolume library (Royer et al., 2015). 3D reconstructions are scaled with gamma adjustment. All imaging experiments were performed at 37 °C, 5 % CO_2_, 20 % O_2_.

### Small molecules

Small molecule concentrations are described in Table S1. All stock solutions were prepared in DMSO. All pharmacological experiments were performed in the presence of 10 µM Verapamil to inhibit drug efflux.

## Supplemental Figure Legends

**Figure S1.**
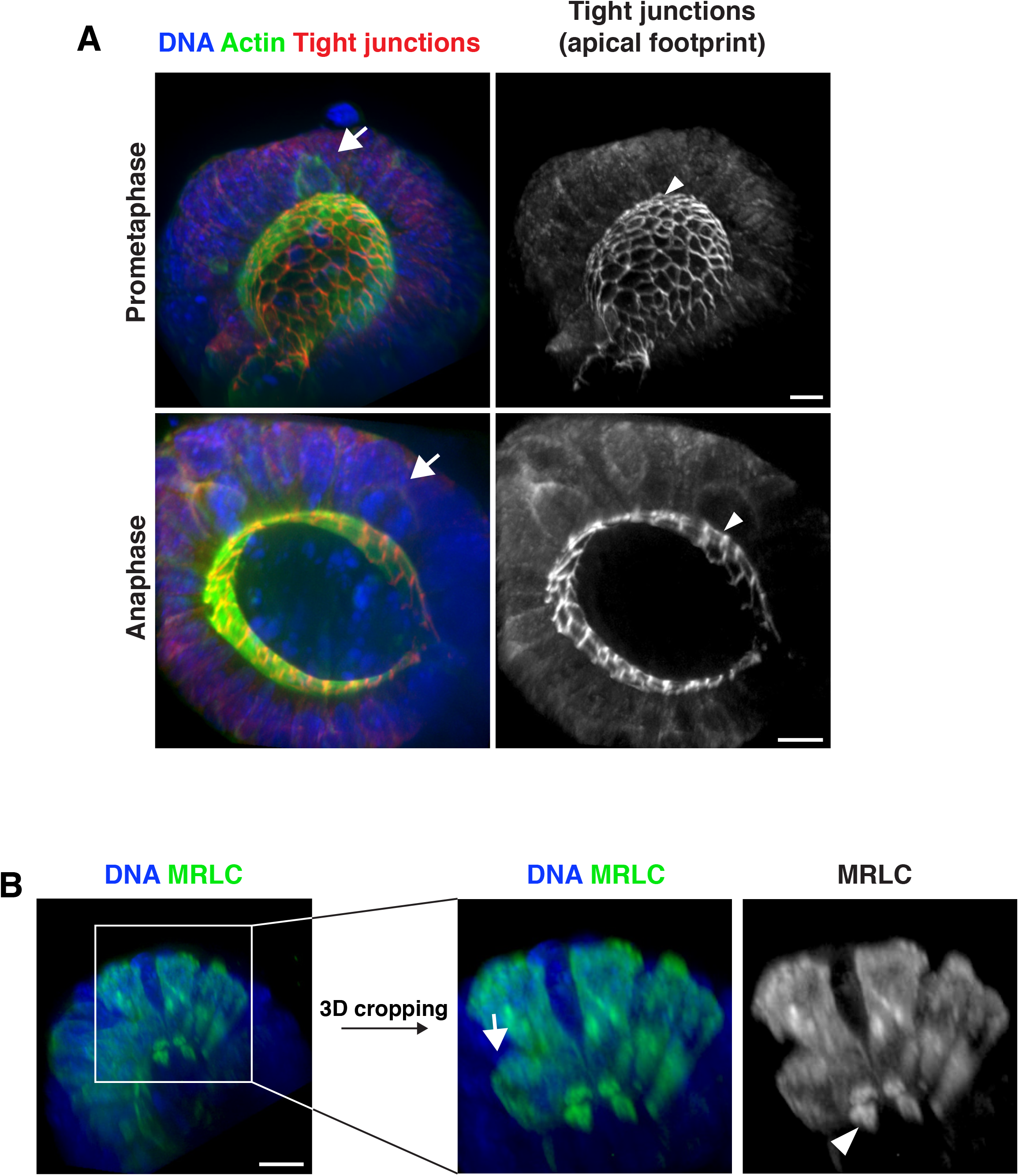
Effects of mitotic progression on the apical surface. A) 3 dimensional reconstruction of immunofluorescence images. Arrow indicates mitotic cell. Arrowhead indicates the corresponding apical footprint. DNA was labeled with Hoechst 33342, actin was labeled with Alexa488-phalloidin and tight junctions were labeled with anti-ZO-1. Top: prometaphase footprint. Bottom: anaphase footprint. B) 3 dimensional reconstruction of immunofluorescence images. Boxed area was cropped in three dimensions to generate right panels. Arrow indicates mitotic cell. Arrowhead indicates the corresponding apical footprint. DNA was labeled with Hoechst 33342. Organoids transduced with MRLC2-mScarlet were used to report on myosin localization. This process can generate mosaic organoids, in which only a subset of cells expresses the transgene, allowing for assignment of the myosin localization to specific cells. Scale bars, 10 µm.

**Figure S2.**
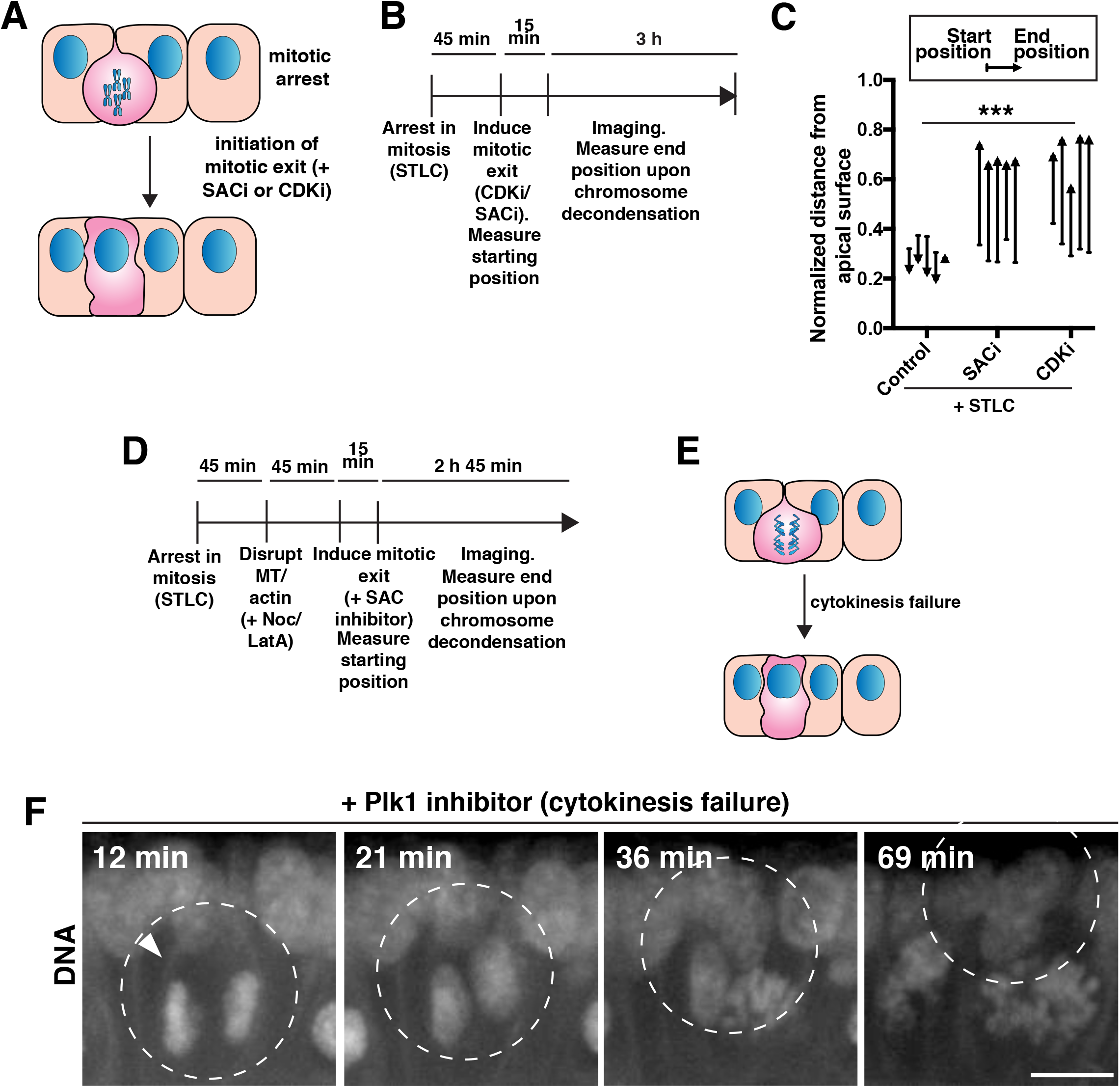
Cytoskeletal requirements for basal reinsertion. A) Cartoon illustrating the strategy used to analyze chromosome movements in organoids following mitotic arrest with S-trityl-L-cysteine (STLC) and subsequent pharmacological disruption of the spindle assembly checkpoint (SAC, using the Mps1 inhibitor AZ3146) or cyclin-dependent kinase (CDK, using the CDK inhibitor RO-3306). B) Schematic of assay used to assess the effects of induced mitotic exit as depicted in (A). C) Quantification of DNA position before the addition of SAC or CDK inhibitors (starting position), and at chromosome decondensation (end position) as depicted in (A) and (B). Each arrow corresponds to the position of the DNA from one cell, before and after the treatment, normalized to the total apical-basal distance of the epithelium. Arrowheads point towards the end position. The end point was defined as chromosome decondensation for the + SAC and + CDK inhibitors conditions. Since the control case does not undergo mitotic exit, the end point was defined as the end of the experiment, after 3 hr of imaging, which is substantially before the organoid begins to die from the treatment. n = 5, ***: p < 0.001, ANOVA of distances moved (end position - start position). D) Schematic of assay used to assess chromosome movements in organoids following mitotic arrest with STLC, pharmacological disruption of the cytoskeleton, and induction of mitotic exit with the SAC inhibitor AZ3146. (Fig. 3B, C, D). E) Cartoon illustrating the strategy used to analyze chromosome movements in organoids following inhibition of cytokinesis with the Plk1 inhibitor BI2536 after mitotic exit. F) Frames from time-lapse imaging of cells in live organoids treated with the Plk1 inhibitor BI2536 to inhibit cytokinesis. DNA was labeled with H2B-mScarlet. Arrowhead indicates cell membrane (labeled with the *R26^mTmG^* allele in the absence of recombination) showing absence of cytokinetic furrow ingression. Time following BI2536 addition is indicated. We note a significant delay in the observation of pharmacological effects on the organoids compared to cultured cells, allowing for initial chromosome alignment and satisfaction of the SAC before the effects of the BI2536 were observed. Representative of n = 8. Scale bar, 10 µm.

**Table S1.** Small molecules used in this study.

| Molecule | Function | Source | Cat # | Final concentration |
| --- | --- | --- | --- | --- |
| Nocodazole | Microtubule inhibitor | Calbiochem | 487929 | 5 μM |
| Latrunculin A | F-actin inhibitor | Calbiochem | 428026 | 4 μM |
| SiR DNA | DNA dye | Cytoskeleton Inc | CY-SC007 | 1 μM |
| Verapamil | Efflux pump inhibitor | Cytoskeleton Inc | CY-SC007 | 10 μM |
| MG132 | Proteasome inhibitor | Sigma | ML449 | 10 μM |
| STLC | Eg5 inhibitor | Sigma | 164739 | 10 μM |
| RO-3306 | CDK inhibitor | Calbiochem | 217699 | 10 μM |
| AZ3146 | Mps1 inhibitor | Tocris | 3994 | 2 μM |
| BI2536 | Plk1 inhibitor | Selleck Biochem | S1109 | 10 μM |
| Tamoxifen | Cre-ER inducer<br>(applied for 6-16 h) | Sigma | T5648 | 1 μM<br>(*Atoh1<sup>CreER</sup>*)<br>0.1 μM<br>(*Vil1<sup>CreER</sup>*) |
| S -- Blebbistatin | Myosin II inhibitor | Abcam | ab120491 | 200 μM |
| S -- Blebbistatin | Myosin II inhibitor | Cayman | 13013 | 200 μM |
| Y27632 | ROCK inhibitor | Selleck | S1049 | 10 μM |

## Supplemental Movie Legends

**Movie S1. Cell behaviors in intestinal organoids.** Membrane-tomato labeled organoids (*R26^mTmG^* in the absence of recombination) imaged with 20X objective at 7 min time points.

**Movie S2. Chromosome movements in intestinal organoids.** H2B-GFP labeled organoids were imaged by SPIM using 40X objectives at 2 min time points.

**Movie S3. Microtubule dynamics during mitosis.** EB3-GFP organoids imaged from early prophase with 60X objective at 20 sec time points.

**Movie S4. Chromosome movements at mitotic onset in latrunculin A-treated organoids.** DNA labeled with SiR-DNA dye imaged with 40X objective at 4 min time points.

**Movie S5. Chromosome movements at mitotic onset in nocodazole-treated organoids.** DNA labeled with SiR-DNA dye imaged with 40X objective at 4 min time points. The cell does not undergo chromosome segregation as it is unable to assemble a mitotic spindle.

**Movie S6. Chromosome movements at mitotic onset in control organoids.** DNA labeled with SiR-DNA dye imaged with 40X objective at 4 min time points.

**Movie S7. Cytokinesis in the intestinal organoids.** MRLC-mScarlet organoids imaged with 60X objective at 20 sec time points.

**Movie S8. Cytokinesis in the intestinal organoids.** MRLC-mGFP organoids imaged down the cytokinetic furrow with 60X objective at 20 sec time points.

**Movie S9. Basal blebbing during cytokinesis.** Rare membrane-GFP cells in organoids generated by stochastic recombination of the *R26^mTmG^* reporter as described in Fig. 1D were imaged with 60x objective at 20 sec time points. Note that the division occurred along the imaging plane, such that the other daughter cell is “behind" the imaged daughter cell.

**Movie S10. Furrow ingression in dissociated intestinal cells.** 3D reconstruction of a single time point from live imaging of mitotic exit of cells dissociated from an EB3-GFP organoid. The reconstruction is rotated away from the viewer. Imaging was performed with a 60X objective. Movie S11. Cell reinsertion behavior. A daughter within the organoid labeled with membrane-GFP (*R26^mTmG^*) blebbing and “crawling" to the basal surface following cytokinesis (note that the division occurred along the imaging plane, such that the other daughter cell is “behind" the imaged daughter cell). Imaged with a 60X objective at 20 sec time points.

**Movie S12. Chromosome movements following induced-mitotic exit in STLC treated organoids.** DNA labeled with SiR-DNA dye imaged with 40X objective at 4 min time points.

**Movie S13. Chromosome movements following induced-mitotic exit in STLC and nocodazole treated organoids.** DNA labeled with SiR-DNA dye imaged with 40X objective at 4 min time points.

**Movie S14. Chromosome movements following induced-mitotic exit in STLC and latrunculin A-treated organoids.** DNA labeled with SiR-DNA dye imaged with 40X objective at 4 min time points.

**Movie S15. Daughter cell separation after division.** Cells of the secretory lineage (red) interspersed with non-secretory cells (green membranes) imaged with 20X objective at 3 min time points.

**Movie S16. 3D reconstruction of separated daughter cells.** Cell of the secretory lineage following mitosis imaged by SPIM with 40X objectives at 2 min time points. Separated daughters are then rotated toward the viewer.

**Movie S17. Insertion of neighbor cell into the cytokinetic furrow.** Cell of the secretory lineage (red) inserts into the furrow of dividing non-secretory cells (green membranes) imaged by SPIM with 40X objectives at 2 min time points. Second clip isolates only the cell of the secretory lineage.

